# TGFβ in the obese liver mediates conversion of NK cells to a less cytotoxic ILC1-like phenotype

**DOI:** 10.1101/576538

**Authors:** Antonia O. Cuff, Francesca Sillito, Simone Dertschnig, Andrew Hall, Tu Vinh Luong, Ronjon Chakraverty, Victoria Male

## Abstract

The liver contains both NK cells and their less cytotoxic relatives, ILC1. Here, we investigate the role of NK cells and ILC1 in the obesity-associated condition, non-alcoholic fatty liver disease (NAFLD). In the livers of mice suffering from NAFLD, NK cells are less able to degranulate, express lower levels of perforin and are less able to kill cancerous target cells than those from healthy animals. This is associated with a decreased ability to kill cancer cells in vivo. On the other hand, we find that perforin-deficient mice suffer from less severe NAFLD, suggesting that this reduction in NK cell cytotoxicity may be protective in the obese liver, albeit at the cost of increased susceptibility to cancer. The decrease in cytotoxicity is associated with a shift towards a transcriptional profile characteristic of ILC1, increased expression of the ILC1-associated proteins CD200R1 and CD49a, and an altered metabolic profile mimicking that of ILC1. We show that the conversion of NK cells to this less cytotoxic phenotype is at least partially mediated by TGFβ, which is expressed at high levels in the obese liver. Finally, we show that reduced cytotoxicity is also a feature of NK cells in the livers of human NAFLD patients.

## Introduction

Natural killer (NK) cells are innate lymphoid cells that recognize and kill virally infected and cancerous cells. In mice, NK cells were long defined as Lineage-negative NK1.1^+^. However, it has recently come to light that, in tissues, cells defined in this way contain at least two distinct populations: circulating, or conventional, NK cells that are CD49a^−^CD49b^+^ and tissue-resident NK-like cells that are CD49a^+^CD49b^−^ (Peng and Tian, 2017). A population of cells that is in many ways equivalent to mouse tissue-resident CD49a^+^ NK cells is also present among CD3^−^CD56^+^ cells in human livers (Male, 2017). These have been called either tissue-resident NK cells or innate lymphoid cells, type 1 (ILC1): here, we call them ILC1 (Peng and Tian, 2015). The finding that Lin^−^ NK1.1^+^ cells in mice and CD3^−^CD56^+^ cells in humans are in fact heterogeneous populations means that it is now necessary to reexamine the roles of these cells, distinguishing between NK and ILC1 subpopulations. This is particularly true in the liver, which contains a large number of ILC1.

Non-alcoholic fatty liver disease (NAFLD) is a spectrum of disease, usually associated with obesity, in which excess fat builds up in the liver. Simple steatosis can progress to a chronic inflammatory condition known as non-alcoholic steatohepatitis (NASH). This in turn progresses to fibrosis and ultimately cirrhosis, which increases the risk of developing hepatocellular carcinoma. NASH patients have increased numbers of CD3^−^CD56^+^CD57^+^ cells, which are likely to represent circulating NK cells, in their livers (Kahraman et al, 2010). The expression of NK cell-activating ligands is also increased in the livers of NASH patients. Recent findings on NK cell dysfunction in obesity (Viel et al, 2017; Tobin et al, 2017; Michelet et al, 2018; O’Shea and Hogan, 2019) and the metabolic activities of NK cells (Viel et al, 2016; Zaiatz-Bittencourt et al, 2018) have led to increased interest in a potential role for NK cells and ILC1 in NAFLD pathogenesis but the field is still in its infancy, not least because most previous studies did not distinguish between NK cells and ILC1 (Luci et al, 2019).

NK cells can limit fibrosis in liver diseases of various etiologies by controlling the activity of hepatic stellate cells (Radaeva et al, 2006; Male et al, 2017). In a mouse study in which a NASH-like disease was induced using a methionine and choline deficient diet, IFNγ production by NKp46^+^ NK cells and ILC1 protected against disease (Tosello-Trampont et al, 2016). On the other hand, NK cells seem to promote obesity, with NK cell-deficient mice gaining less weight in response to obesogenic diets (Wensveen et al, 2015; Lee et al, 2016; O’Rourke et al, 2014) and displaying less fat accumulation and inflammation in their livers (Cepero-Donates et al, 2016). There is also evidence that NK cells can be harmful in chronic inflammatory liver disease, where they contribute to pathology by killing hepatocytes (Dunn et al, 2007; Beraza et al, 2009). In support of this being the case in NASH, mice lacking either TRAIL (an apoptosis-inducing TNF family ligand expressed by NK cells) or its receptor are partially protected against obesity-associated NASH (Idrissova et al, 2014; Hirsova et al, 2017).

Here, we investigate the activity of NK cells and ILC1 in a mouse model of NASH and in the livers of NAFLD patients. We report that NK cells in the livers of obese mice are less able to degranulate and kill cancerous target cells than those from lean mice, a phenotype that extends to a reduced in vivo ability to kill cancer cells. This decrease in cytotoxicity is associated with a shift towards an ILC1-like phenotype, which seems to be at least partially mediated by high levels of TGFβ produced in the obese liver. Finally, we show that in humans, as in mice, NK cells from obese livers are less able to degranulate and kill.

## Results

### NK cells in the livers of obese mice are less cytotoxic than those in the livers of lean mice

To investigate the activity of NK cells and ILC1 in the liver during obesity-associated liver disease, we examined the spleens and livers of mice that were kept for up to 24 weeks on a high fat and sugar diet (Oben et al, 2010). As previously reported, mice on the diet became obese (Figure 1a), accumulated fat in their livers (Figure 1b) and displayed dysregulated glucose homeostasis (Figure 1c). Mice also displayed histological evidence of NAFLD (Figure 1d,e) and increased circulating alanine transaminase (ALT), which is an indicator of liver damage (Figure 1F).

**Fig. 1.**
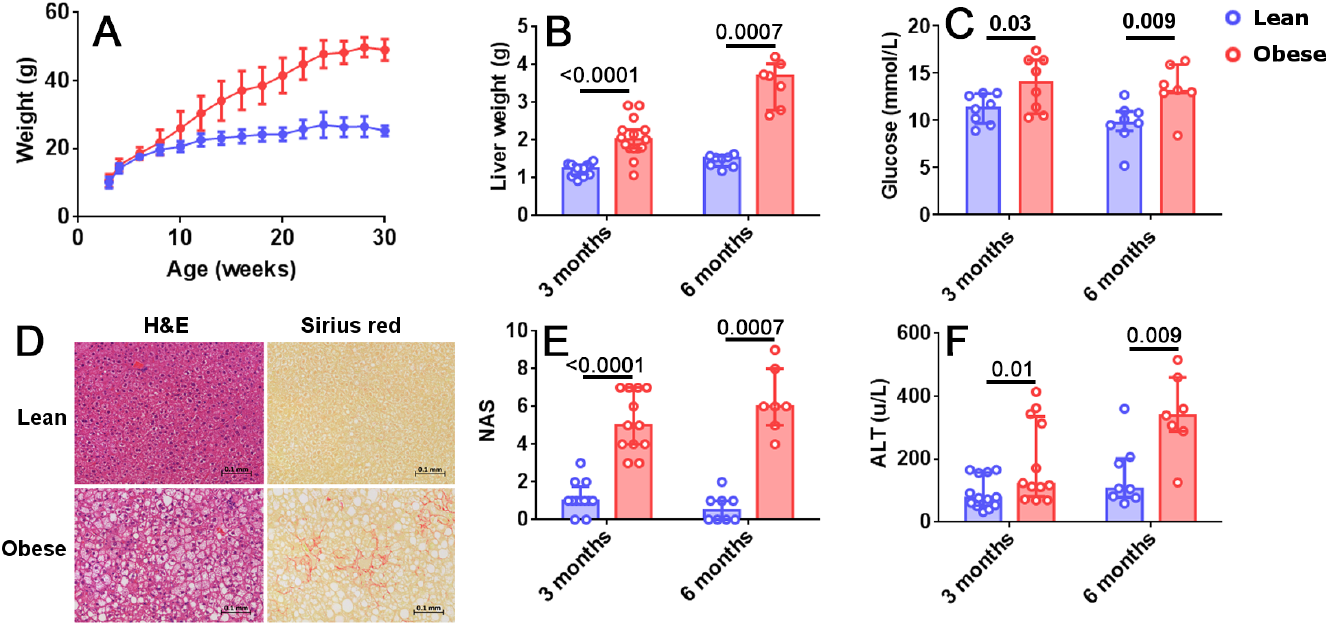
Mice fed an obesogenic diet develop fatty liver disease. A. Growth curves for mice fed an obesogenic diet (red) or standard chow (blue). B. Liver weights at 3 (n = 12 mice/group) or 6 months (n = 8 mice in the lean and 7 mice in the obese group). C. Plasma glucose levels (n = 8 mice per group, except for obese at 6 months where n = 7 mice). D. H and E and Picrosirius red staining in representative livers after 3 months on the obesogenic or standard diet. E. Histological scores (n = 12 mice per group at 3 months, 8 lean mice at 6 months and 7 obese mice at 6 months). F. Plasma ALT levels (n = 12 mice per group at 3 months, 8 lean mice at 6 months and 7 obese mice at 6 months). Significance was determined using Mann Whitney U Tests; medians and IQRs are shown.

NK cells (defined as Lineage-negative NK1.1^+^CD49a^−^CD49b^+^) isolated from the livers of mice that had been kept on the obesogenic diet for 12 weeks degranulated less than those from the livers of their lean littermates (Figure 2a,b). This was also the case for NK cells isolated from spleens, although the reduction was smaller (a difference in the medians of 4.3% in splenic NK cells, compared to 10.0% in liver NK cells; Figure 2b). We observed no difference in the degranulation of liver ILC1 (defined as Lineage-negative NK1.1^+^CD49a^+^CD49b^−^) between lean and obese mice. We also found a significant reduction in the expression of perforin by NK cells in the livers of obese mice, that we did not detect in splenic NK cells (Figure 2c). This suggests that NK cells from the livers of obese mice are both less able to degranulate and less able to kill target cells than those from their lean littermates. We did not observe any difference in the expression of granzyme B (Figure 2d) although this may be accounted for by the low levels at which this protein is known to be expressed in unstimulated mouse NK cells (Fehniger et al, 2007).

**Fig. 2.**
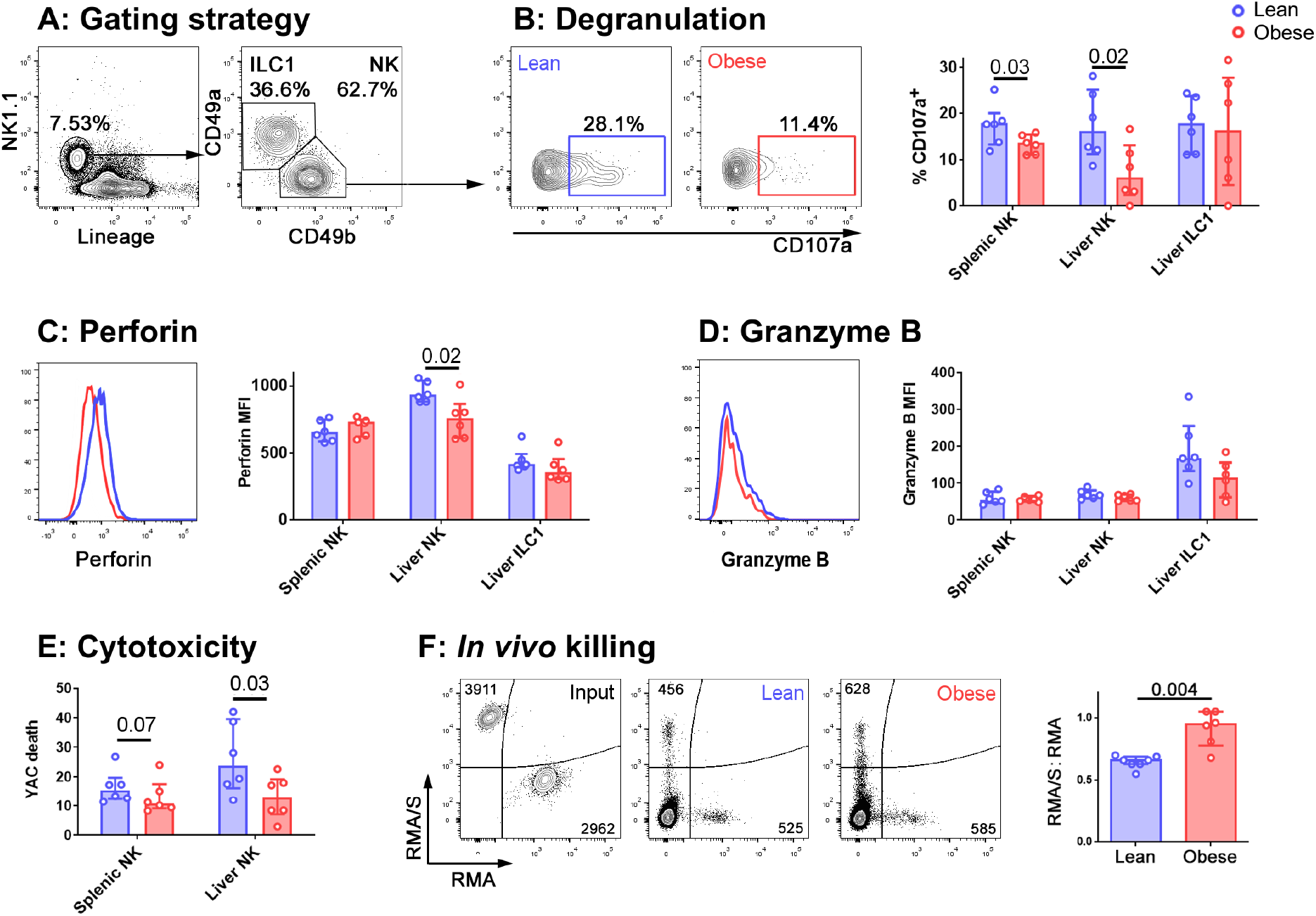
NK cells in the livers of obese mice are less cytotoxic than those in lean mice. A. Immune cells were isolated from mouse livers. NK cells were identified by scatter, and as live CD45^+^ Lineage-negative NK1.1^+^ CD49b^+^ cells. ILC1 were identified as live CD45^+^ Lineage-negative NK1.1^+^ CD49a^+^ cells. B. Intrahepatic leukocytes were cultured for 4h in the presence of anti-CD107a and Brefeldin A. Representative CD107a staining of NK cells from a lean (left, blue) and an obese (right, red) mouse and summary data are shown. C. Representative perforin staining in liver NK from a lean (blue trace) and an obese (red trace) mouse. MFI of perforin in splenic NK, liver NK and liver ILC1 from lean and obese mice are shown. D. Representative granzyme B staining in liver NK from a lean (blue trace) and an obese (red trace) mouse. MFI of granzyme B in splenic NK, liver NK and liver ILC1 from lean and obese mice are shown. E. NK cells were sorted from the spleens and livers of lean and obese mice and cultured with the cancerous NK cell target line YAC-1. YAC-1 death at 24h is shown. F. CTV-labelled RMA/S (NK cell targets) CTB-labelled RMA (recovery control) cells were mixed in a 1:1 ratio and intravenously injected into lean or obese mice. Representative input cells (leftmost, black) and recovered cells from lean (center, blue) and obese (rightmost, red) mice are shown. The RMA/S:RMA ratio in recovered splenocytes, normalised to input ratio, is shown. For all panels, n = 6 mice per group; significance was determined using Mann Whitney U Tests; medians and IQRs are shown.

The reduced ability of NK cells in obese mice to degranulate suggests that their ability to kill cancerous target cells will be impaired. To determine whether this was the case, and to examine the relative defects in NK cell killing in the spleens and livers of obese compared to lean mice, we sorted NK cells from the two organs and assessed their ability to kill YAC-1 cancerous target cells in vitro. NK cells from both the spleens and livers of obese mice were less able to kill YAC-1 cells than those from their lean littermates, although the defect in killing was only significant (at p < 0.05) in NK cells isolated from the liver (Figure 2e). Similar to our observations of degranulation, the reduction in target cell killing was greater in NK cells isolated from livers than those isolated from spleens (a difference in the medians of 2.7% in splenic NK cells, compared to 10.1% in liver NK cells).

To determine whether this affects target cell killing in vivo, we next injected mice intravenously with equal numbers of fluorescently-labelled RMA/s cells (which are MHC class I-negative NK targets) and RMA cells (which are not NK targets and act as a recovery control) (Kärre et al, 1986). We collected spleens from the recipient mice and determined the number of cells remaining in each population by flow cytometry. The lean mice had eliminated the RMA/s cells with approximately 40% efficiency. In contrast, the obese mice had eliminated them with only 5% efficiency (Figure 2f).

### Perforin-mediated cytotoxicity promotes fibrosis in NAFLD

Clearly, the diminished ability of NK cells to kill target cells in obese animals is likely to limit cancer surveillance. However, since we observed the largest reductions in both degranulation and cytotoxicity in NK cells isolated from the liver, and since the reduction in perforin expression seemed to particularly affect NK cells in the liver, we sought to define the likely effect of these changes in NAFLD. Given that TRAIL-mediated cytotoxicity exacerbates NAFLD (Idrissova et al, 2014; Hirsova et al, 2017), we speculated that there might be some advantage to the reduced perforin-mediated cytotoxicity we observed in the obese liver. We hypothesized that perforin-mediated killing is harmful in the liver during obesity, and that the reduction in NK cell degranulation, perforin expression and cytotoxicity could therefore be protective. In this case, we would expect mice lacking perforin to suffer less severe liver disease in obesity. We therefore determined the extent of liver damage in wild type and perforin-deficient mice that had been kept on the obesogenic diet for 24 weeks. This later timepoint was chosen so that we could compare the development of fibrosis, which is not usually pronounced at 12 weeks in this model, between the two genotypes.

There was no difference in weight gain (Figure 3a), hepatomegaly (Figure 3b) or glucose homeostasis (Figure 3c) between the wild type and perforin knockout mice. Therefore, unlike mice that completely lack NK cells, perforin-deficient mice gain weight and alter their glucose homeostasis similarly to wild type mice (Wensveen et al, 2015; Lee et al, 2016; O’Rourke et al, 2014). This suggests that any differences we observe in liver pathology are unlikely to occur as a result of less severe metabolic dysregulation in these mice. We observed a significantly lower histopathological score in the perforin-deficient mice than the wild type controls (Figure 3d,e) and decreased circulating ALT, although this was not significant at p < 0.05 (Figure 3f). Fibrosis, determined by Picrosirius red staining, was significantly lower in perforin-deficient mice (Figure 3d,g) and real time PCR for markers of fibrosis revealed that the livers of the perforin-deficient mice expressed less Acta2 (a smooth muscle actin; Figure 3h) and Col1a1 (collagen type I alpha 1; Figure 3i) than those of controls, although the difference was only significant at p < 0.05 for Acta2.

**Fig. 3.**
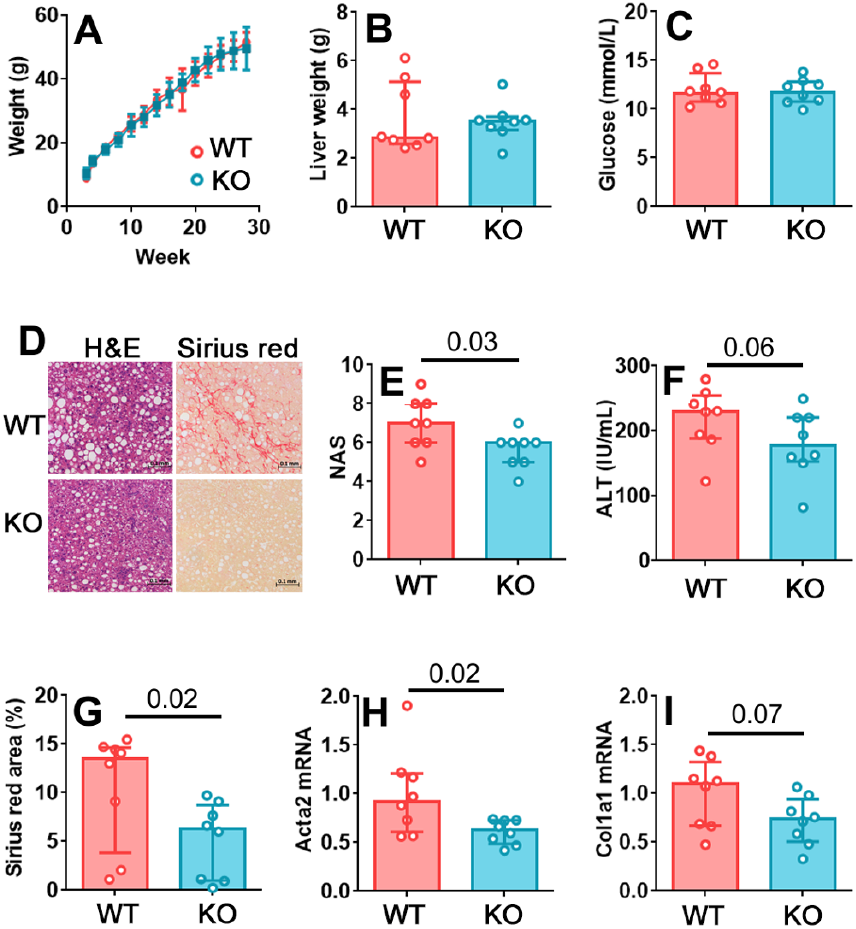
Perforin-mediated killing promotes fibrosis in NAFLD. A. Growth curves for wild type and perforin knockout mice on the obesogenic diet. B. Liver weights for wild type and perforin knockout mice after 24 weeks on the obesogenic diet. C. Plasma glucose levels. D. Representative H and E and Picrosirius red staining in a wild type and a perforin knockout mouse. E. Histological score in wild type and perforin knockout mouse livers, after 24 weeks on the obesogenic diet. F. Plasma ALT levels. G. Picrosirius red positive area in representative histological fields. H and I. Acta2 and Col1a1 mRNA in livers of wild type and perforin knockout mice, after 24 weeks on the obesogenic diet. n = 8 mice per group; significance was determined using Mann Whitney U Tests; medians and IQRs are shown.

### NK cells in the livers of obese mice display a transcriptional profile characteristic of ILC1

We next sought to define the mechanism by which the reduction in NK cell cytotoxicity occurs. To achieve this, we analyzed the transcriptomes of NK cells and ILC1 in the livers of lean and obese mice by RNASeq. Raw RNAseq data and differentially expressed gene lists are available from the National Center for Biotechnology Information Gene Expression Omnibus under accession no. GSE122828.

18 of the 20 genes identified as being significantly overexpressed (FC > 2, p_adj_ < 0.05) in NK cells from obese mice compared to those from their lean littermates, were also overexpressed in ILC1 compared to NK cells (Figure 4a; genes characteristic of ILC1 are marked with an asterisk). We particularly noted that transcripts encoding the inhibitory receptors CD200R1, LAG3 and CD101, all of which are involved in limiting immune pathology in inflammatory diseases (Hoek et al, 2000; Fernandez et al, 2007; Snelgrove et al, 2008; Anderson et al, 2016) were increased in NK cells from obese mice compared to lean, as well as in ILC1 compared to NK cells. We confirmed the increase in expression at the protein level for CD200R1 (Figure 4b) and LAG3 (Figure 4c).

**Fig. 4.**
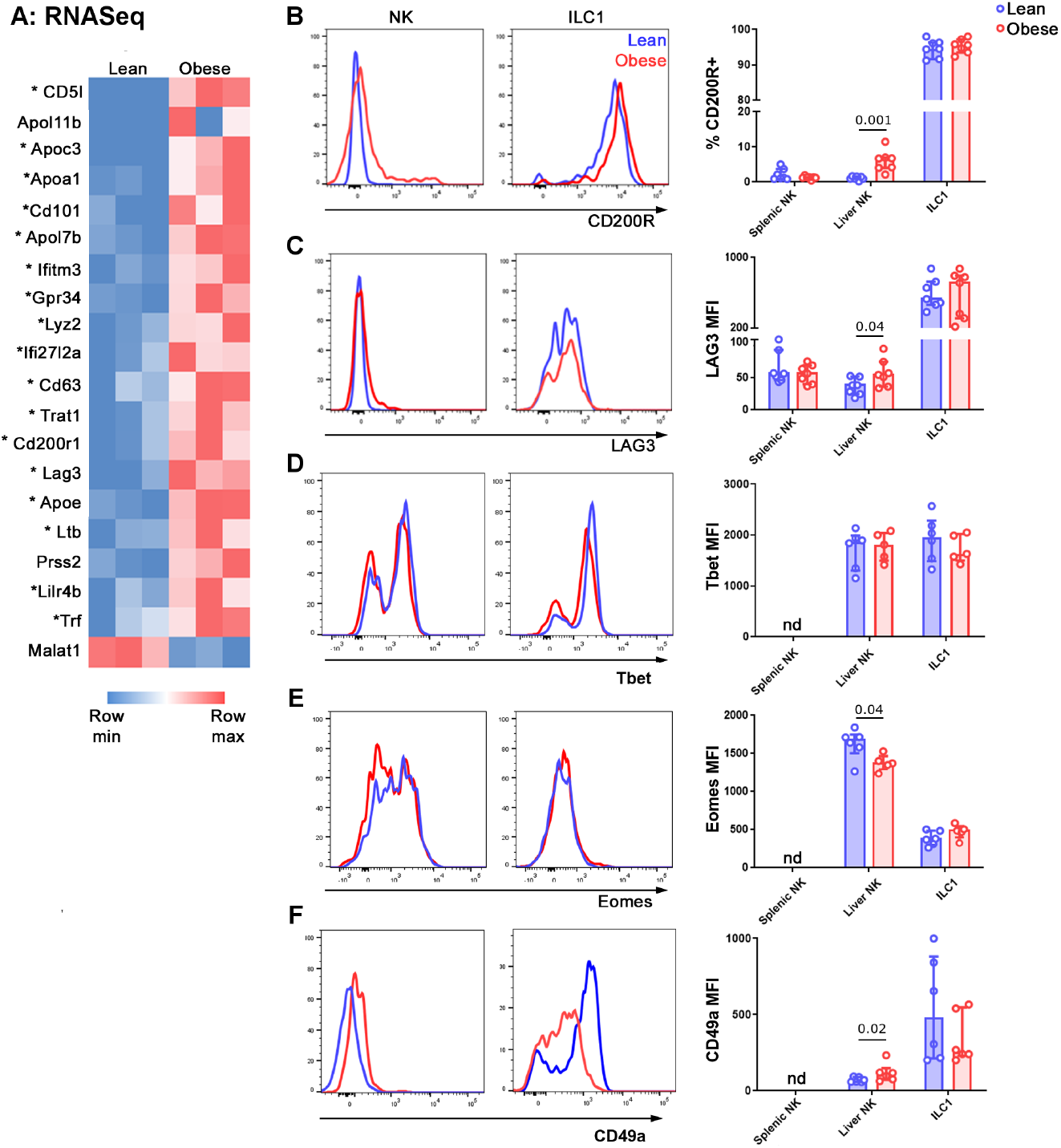
NK cells in the livers of obese mice acquire an ILC1-like transcriptional profile. A. NK cells and ILC1 were sorted from the livers of lean and obese mice and examined by RNASeq. Genes that were differentially expressed by > 2-fold, with p_adj_ < 0.05, in NK cells of lean versus obese mice are shown. Genes that are also differentially expressed in ILC1 compared to NK (> 2-fold, with p_adj_ < 0.05) are marked with an asterisk. B,C. Protein expression of the inhibitory receptors CD200R1 (B) and LAG3 (C) in NK cells and ILC1 of lean (blue) compared to obese (red) mice. D-E. Expression of the ILC1-associated proteins Tbet (D), Eomes (E) and CD49a (F) in NK cells and ILC1 of lean (blue) compared to obese (red) mice. For panel F, NK cells were defined as Lineage-negative NK1.1^+^ CD49b^+^ and ILC1 were defined as Lineage-negative NK1.1^+^ CD49b^−^. n = 6 mice per group; significance was determined using Mann Whitney U Tests; medians and IQRs are shown.

NK cells and ILC1 express the transcription factor Tbet at roughly similar levels, although they rely on it to different extents (Sojka et al, 2014). Consistent with this, we did not detect any change in the expression of Tbet in NK cells of the livers of lean compared to obese mice (Figure 4d). On the other hand, the transcription factor Eomes is expressed only by NK cells and specifies the NK cell lineage (Daussy et al, 2014). We observed decreased expression of Eomes in NK cells in the livers of lean compared to obese mice (Figure 4e). We also examined the expression of the ILC1-associated integrin CD49a and found a small but significant increase in its expression by NK cells in the livers of obese compared to lean animals (Figure 4f), consistent with a shift towards an ILC1-like phenotype.

### NK cells in the livers of obese mice have an altered metabolic profile

Gene ontogeny analysis on the differentially expressed transcripts (Figure 4a) further revealed that the most overrepresented pathway in the NK cells of obese mice, compared to those of lean littermates, was Triglyceride catabolic processes (p = 3.65 x 10-3). We therefore examined the cells for signs of altered immunometabolism.

Scatter has been used as an indicator of metabolic alteration in NK cells (Castro et al, 2018) and we found that side scatter increased in NK cells from the livers of obese mice (Figure 5a). NK cells use mTOR for nutrient sensing and it also has an important role in immune regulation (O’Brien and Finlay, 2019). The phosphorylation of ribosomal S6 protein (pS6) is used as an indicator of mTORC1 signaling and we found that pS6 was increased in NK cells freshly isolated from the livers of obese compared to lean mice (Figure 5b). Notably, in both these cases, the metabolic alterations that we observed in the NK cells served to make them more ILC1-like, similar to our findings at the transcriptional level. We also observed an increase in the expression of the glucose transporter Glut1 (Figure 5c).

**Fig. 5.**
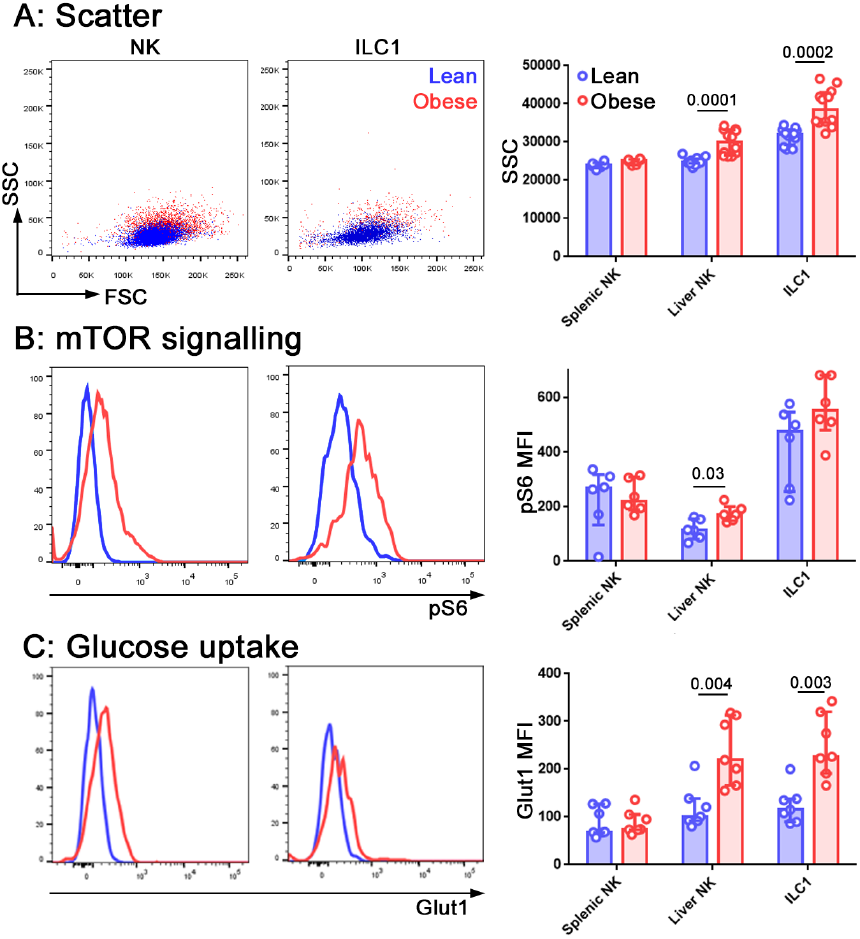
NK cells in the livers of obese mice are metabolically reprogrammed. Forward and side scatter (A), pS6 staining (B) and Glut1 staining (C) of freshly isolated NK cells and ILC1 from lean (blue) and obese (red) mice. n = 12 mice per group (A), 6 mice per group (B) and 7 mice per group (C); significance was determined using Mann Whitney U Tests; medians and IQRs are shown.

### TGFβ in the obese liver limits NK cell degranulation and alters their metabolic profile

The conversion of NK cells to a less cytotoxic ILC1-like phenotype has previously been reported in a number of situations and in all these cases, TGFβ was a key cytokine driving the conversion (Cuff et al, 2016; Gao et al, 2017; Cortez et al, 2017). We therefore hypothesized that TGFβ in the obese liver might be mediating the reduction in cytotoxic ability and the acquisition of ILC1-like features that we observed. Tgfb1 transcript was increased in the livers of obese compared to lean mice (Figure 6a) and we also observed increased TGFβ1 protein in conditioned medium from obese compared to lean livers (Figure 6b) and in the plasma of obese mice (Figure 6c).

**Fig. 6.**
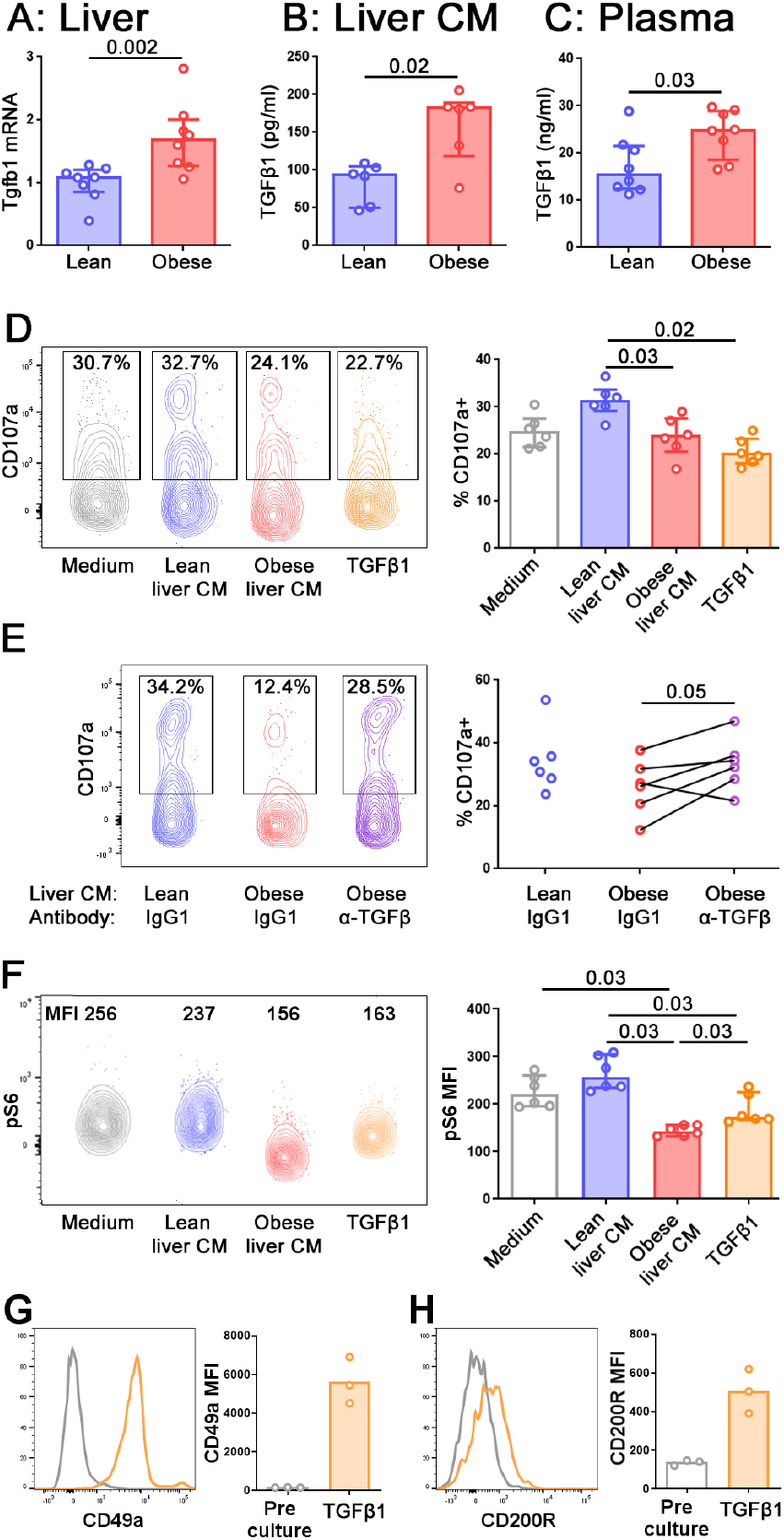
TGFβ in obese mouse livers limits NK cell degranulation and alters their metabolic profile. A. Tgfb1 mRNA in the livers of lean and obese mice, normalised so that mean Tgfb1 transcript expression in the livers of lean mice = 1. B,C. TGFβ1 in lean and obese liver conditioned medium (B) and lean and obese mouse plasma (C) measured by ELISA. n = 8 mice per group (A, C) and 6 conditioned media per group (B); significance was determined using Mann Whitney U Tests; medians and IQRs are shown. D. Splenic NK cells were cultured for 24h in medium alone, lean or obese liver conditioned medium, or 10 ng/ml TGFβ1. For the last 4h of culture, CD107a antibody and Brefeldin A were added to media. n = 6 culture conditions per group; significance was determined using Mann Whitney U Tests with Holm’s correction for multiple comparisons; medians and IQRs are shown. E. Splenic NK cells were cultured as described in D, with the addition of 10 μg/ml anti-TGFβ antibody or isotype control. n = 6 culture conditions per group; significance was determined using a Wilcoxon signed ranks test, where each pair is obese liver conditioned medium from the same mouse, with anti-TGFβ antibody or isotype control. Degranulation in NK cells cultured in lean liver conditioned medium with isotype control are shown for comparison, but were not statistically tested. F. Splenic NK cells were cultured for 24h in medium alone, lean or obese liver conditioned medium, or 10 ng/ml TGFβ1, before staining for pS6. n = 6 culture conditions per group; significance was determined using Mann Whitney U Tests with Holm’s correction for multiple comparisons; medians and IQRs are shown. G, H. Splenic NK cells were cultured for 5d in 10 ng/ml TGFβ1. CD49a (G) and CD200R (H) staining before and after culture are shown. For this small dataset, no statistical testing was carried out.

When NK cells from the spleens of lean animals were cultured for 24 hours in conditioned medium from either lean or obese mouse livers, we observed that their ability to degranulate was impaired following culture in obese, compared to lean, liver conditioned medium. We were able to recapitulate this phenotype by culturing the NK cells in recombinant TGFβ1 (Figure 6d). Furthermore, the addition of an anti-pan TGFβ antibody to the cultures partially rescued the ability of the NK cells to degranulate following culture with obese liver conditioned medium (Figure 6e), suggesting that TGFβ in the obese liver limits the ability of NK cells to degranulate. We also examined pS6 expression as an indicator of mTORC signaling. Consistent with previous reports, NK cells cultured with TGFβ1 displayed reduced pS6 (Viel et al, 2016), as did NK cells cultured in obese liver conditioned medium (Figure 6f), suggesting that TGFβ in the obese liver can also affect the metabolic profile of NK cells.

Consistent with previous reports (Gao et al, 2017), NK cells cultured for 5 days in TGFβ1 upregulated ILC1-associated molecules such as CD49a (Figure 6g) and CD200R1 (Figure 6h). However, we did not observe this phenomenon in NK cells cultured for only 24 hours in TGFβ1 and we were unable to keep NK cells alive in liver conditioned medium for 5 days. Therefore, we were unable to directly assess the ability of liver conditioned medium to convert NK cells to ILC1. Our findings that the obese liver contains high levels of TGFβ1 and that TGFβ can convert NK cells to an ILC1-like phenotype does, however, suggest that this could occur in vivo.

### NK cells in the livers of NAFLD patients are less cytotoxic than those from healthy controls

Our observations in mice prompted us to ask whether NK cells in the livers of NAFLD and NASH patients are also less able to degranulate than those from healthy controls. We isolated intrahepatic leukocytes from seven healthy livers, six livers with simple steatosis (livers that showed steatosis but no inflammation) and six livers displaying various levels of inflammation and fibrosis in addition to steatosis. We found the ability of NK cells (defined as CD3^−^CD56^+^Tbet^hi^ Eomes^lo^; Cuff, 2016, JI; Figure 7a) to degranulate was inversely correlated with disease severity (Figure 7b-d). In seven samples where we were also able to sort the NK cells and assess their ability to kill the classical human NK cell target line K562, we found decreased killing by NK cells from NAFLD livers, compared to healthy controls although we were not able to collect enough data points to assess the statistical significance of this relationship (Figure 7e).

**Fig. 7.**
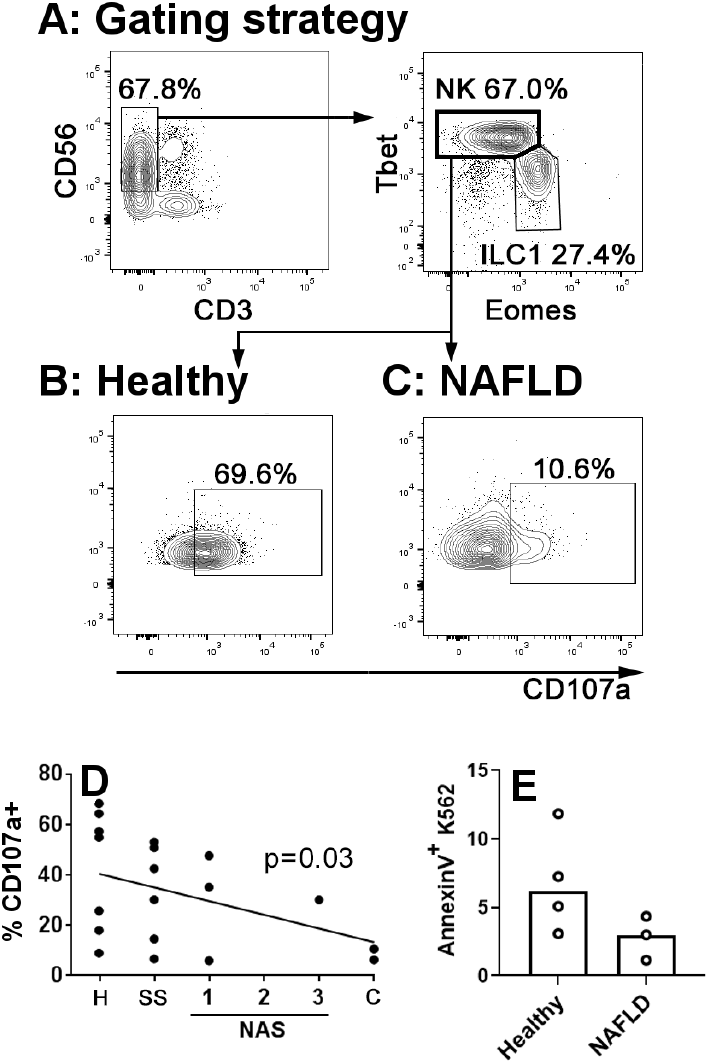
NK cells in the livers of NAFLD patients are less cytotoxic than those from healthy controls. A. Immune cells were isolated from human livers. NK cells were identified by scatter, and as live CD45^+^ CD56^+^ CD3^−^ Tbet^hi^ Eomes^lo^ cells. B and C. Intrahepatic leukocytes were cultured for 4h in the presence of anti-CD107a and Brefeldin A. Representative CD107a staining of NK cells from a healthy (B) and a NAFLD (C) liver is shown. D. Inverse correlation between CD107a staining and histological score in human livers. H: healthy; SS: simple steatosis; NAS: NASH Activity Score; C: cirrhosis. Significance was determined using Spearman’s Rank Correlation. E. CD56^+^ CD3^−^ CD16^+^ CXCR6^−^ NK cells were sorted from healthy (n = 4) and NAFLD (n = 3) livers and cultured for 4h with the human NK target cell, K562. At the end of culture, target cell death was assessed using AnnexinV staining. Medians are shown. For this small dataset, no statistical testing was carried out.

## Discussion

It is becoming increasingly apparent that NK cells are dysfunctional in obesity and that this could be one mechanism that accounts for the well-established link between obesity and cancer (Viel et al, 2017; Tobin et al, 2017; Michelet et al, 2018; O’Shea and Hogan, 2019). These studies have largely focused on NK cells in the blood, but in this study we looked at how NK cells in the liver change during obesity, and the impact that this may have not only on cancer immunosurveillance, but also on NAFLD pathogenesis.

We found that in obesity NK cells in the livers of both humans and mice are less able to degranulate and kill target cells. This is consistent with reports that NK cells taken from the blood of obese humans (Viel et al, 2017; O’Shea et al, 2010; Tobin et al, 2017; Michelet et al, 2018) or the spleens of obese mice (Michelet et al, 2018) are less cytotoxic than those taken from their lean counterparts. We further show that the NK cells of obese mice are less able to kill cancerous target cells in vivo. It has recently been reported that NK cells that had been cultured in fatty acids before being adoptively transferred into a B16 melanoma-bearing host are less able to control the cancer than control NK cells. Obese mice are also less able to control B16 metastases than their lean counterparts, a phenotype that was associated with reduced NK cell infiltration into metastatic foci (Michelet et al, 2018). Our finding that obese mice are less able to control RMA/S cells confirms these findings using another type of cancerous target.

This reduction in NK cell cytotoxicity in obesity is detrimental to tumor immunosurveillance, but one key question is whether it is helpful or harmful in the context of NAFLD. We observed greater reductions in degranulation, perforin expression and cytotoxicity in the liver than in the spleen. Further, TRAIL-mediated cytotoxicity is known to promote pathogenesis in NAFLD (Idrissova et al, 2014; Hirsova et al, 2017). Therefore, we postulated that the decrease in degranulation and perforin-mediated cytotoxicity that we observed in the obese liver might protect against liver disease. In support of this idea, perforin-deficient mice suffered from less severe NAFLD. Since these animals lack perforin globally, we cannot say with certainty that the protection is not mediated at least in part by defective T cell cytotoxicity. Indeed if, as we suggest, the liver is acting to protect itself against immunopathology, it is likely that T cell cytotoxicity will also be reduced. Nevertheless, the protection from liver disease that we find in the absence of perforin does suggest that the defect in NK cell degranulation we observe in the obese liver may act to protect it.

We found the genes differentially expressed between NK cells from lean versus obese livers to be strikingly similar to those that are differentially expressed between NK cells and ILC1, suggesting NK cell conversion to an ILC1-like phenotype in the obese liver. Consistent with this, we found the ILC1-associated proteins CD200R1 (Weizman et al, 2017), LAG3 and CD49a increased in NK cells from obese livers, and expression of the NK cell lineage-defining transcription factor Eomes decreased. NK cells in the livers of obese mice also displayed an altered metabolic profile, which was somewhat similar to that of ILC1.

There are a number of situations in which NK cells are converted to ILC1-like cells and in some of these, notably in the tumor microenvironment, this is associated with a decrease in cytotoxicity similar to our findings in the obese liver (Cuff et al, 2016; Cortez et al, 2017; Gao et al, 2017). NK to ILC1 conversion in all these situations is mediated by TGFβ. Further, TGFβ is known to alter the metabolic profile of NK cells and limit their cytotoxicity (Viel et al, 2017; Zaiatz-Bittencourt et al, 2018). Therefore, we postulated that NK cell acquisition of an ILC1-like phenotype in the obese liver might be mediated by TGFβ. TGFβ1 levels were higher in obese than lean livers, consistent with previous reports in both mice and humans (Yadav et al, 2011; Mouralidarane et al, 2013; Hart et al, 2017). Conditioned medium from obese livers limited the ability of NK cells to degranulate, mirroring the phenotype observed in the obese liver, and this effect was TGFβ-dependent. We also assessed the effect of liver conditioned medium on mTORC1 signaling, as determined by pS6 levels, and again observed obese liver conditioned medium mimicking the effect of TGFβ1 (Viel et al, 2016).

Interestingly, we observed reduced pS6 levels on short term culture with either obese liver conditioned medium or TGFβ1 but increased pS6 levels in the livers of mice that had been kept on the obesogenic diet for six months. It has previously been suggested that there may be differences in the kinetics of altered mTORC1 signaling in short-term versus longer-term TGFβ exposure, and this may be at play here (Zaiatz-Bittencourt et al, 2018). In support of this idea, the short- and long-term effects of obesity on NK cell metabolism are known to differ: in a pediatric cohort, obesity was associated with increased Glut1 and pS6 expression (Tobin et al, 2017), whereas in adults, obesity was associated with decreased 2-NBDG uptake (a proxy for glucose uptake) and pS6 expression (Michelet et al, 2018). It is also likely to be the case that other factors present in the obese liver, for example high levels of fatty acids, affect NK cell cytotoxicity and metabolism (Michelet et al, 2018) and the effects of these may interact with those of TGFβ, possibly in a manner which changes over time. This could also explain our observation that obese liver conditioned medium depresses NK cell pS6 expression more than TGFβ1 alone.

In this study, we have shown that NK cells in the liver display a less cytotoxic ILC1-like phenotype in the context of obesity and that this is likely to be at least partially TGFβ-mediated. Our finding that perforin-mediated killing is harmful in NAFLD suggests that this reduction in NK cell cytotoxicity is protective in the liver, albeit at the cost of increased susceptibility to cancer. As well as mediating conversion of NK cells to ILC1 (Cuff et al, 2016; Cortez et al, 2017; Gao et al, 2017), TGFβ also alters NK cell metabolism and cytotoxicity (Viel et al, 2017; Zaiatz-Bittencourt et al, 2018). It has recently been suggested that NK cell metabolism could be targeted to prevent cancer (O’Brien and Finlay, 2019). In the light of recent findings that NK cells are less cytotoxic in obesity and that this is at least partially mediated by metabolic changes (Viel et al, 2017; O’Shea et al, 2010; Tobin et al, 2017; Michelet et al, 2018), it might be tempting to consider using such treatments as a way to break the well-established link between cancer and obesity (Park et al, 2014). However, the implication that reduced NK cell cytotoxicity may protect the liver during obesity sounds a note of caution about such approaches, which may result in adverse effects in the liver. Indeed, NK cell-derived ILC1-like cells play a role in tissue repair (Gao et al, 2017) so the possibility that the ILC1-like cells we see emerging from NK cells in the obese liver may even have a reparative effect is worthy of further investigation.

## Materials and Methods

### Human liver biopsies

Liver biopsies were taken from livers that were destined for transplantation, but that were discarded because of long warm ischemic time, vascular abnormalities, or tumors elsewhere in the patient (n = 7), or because they displayed signs of NAFLD or NASH on histopathologic examination (n = 10). Biopsies were also taken from livers explanted during transplantation for NASH (n = 2). Intrahepatic leukocytes were isolated as previously described (Cuff et al, 2016). Briefly, liver tissue was finely minced using scalpels, passed through a 70-μm strainer, and the collected cells were layered onto Ficoll (GE Healthcare, Amersham, U.K), centrifuged (400 × g, 20 min, 20°C, light braking), and the interface was taken and washed twice with PBS (750 × g, 15 min, 20°C). Ethical approval for use of human liver samples was obtained through the Royal Free Hospital Biobank (National Health Service Research Ethics Committee approval no. 11/WA/0077, study no. 9455).

### Mice

Female C57BL/6J (bred in house) or Prf1−/− mice (Charles River; RRID MGI:5576721) were randomized at weaning onto standard chow (RM1) or a highly palatable obesogenic diet consisting of 22.6% fat, 23.0% protein and 40.2% carbohydrate (w/w); diet code 824018 – ‘45% AFE fat’ Special Dietary Services, Essex, UK), supplemented with sweetened condensed milk (Nestle) ad libitum (Oben et al, 2010).

After 12 or 24 weeks on the diet, mice were sacrificed by direct cervical dislocation. Blood was collected by cardiac puncture and centrifuged at 10,000 xg for 10 minutes at room temperature to isolate serum for ALT and glucose measurement and pieces of liver were fixed in 10% neutral buffered formalin, fixed, sectioned and stained with H and E or Picrosirius red. Sections were blinded and scored by a hepatopathologist, using the Brunt-Kleiner NASH activity score (NAS; Brunt et al, 2011). Spleens were collected and splenocytes isolated as previously described (Cuff and Male, 2017). Intrahepatic lymphocytes were isolated using an adaptation of the method from Cuff and Male, 2017. Briefly, finely minced liver tissue was collected in RPMI 1640 medium (Life Technologies brand; Thermo Fisher Scientific, Hudson, NH) and passed through a 70 μm cell strainer. The suspension was spun down (500 × g, 4°C, 10 minutes) and the pellet resuspended in RPMI 1640 medium. The cell suspension was layered over 24% Optiprep (Sigma-Aldrich) and centrifuged without braking (700 × g, RT, 20 minutes). The interface layer was taken and washed in HBSS without Ca^2+^ Mg^2+^ (Lonza, distributed by VWR, Lutterworth, UK) supplemented with 0.25% bovine serum albumin (Sigma-Aldrich, Hammerhill, U.K) and 0.001% DNase I (Roche, distributed by Sigma-Aldrich).

Animal husbandry and experimental procedures were performed according to UK Home Office regulations and institute guidelines, under project license 70/8530.

### In vivo NK cytotoxicity assays

RMA and RMA/s thymoma cells (cultured in Iscove’s modified Dulbecco’s medium supplemented with 10% FCS; Life Technologies) were labeled with Cell Trace Blue or Cell Trace Violet, respectively, according to the manufacturer’s instructions (Thermo Fisher Scientific). 10^7^ cells of each cell type were injected intravenously into control or obese mice. After 4h, recipients were sacrificed by direct cervical dislocation and the spleens were examined for fluorescent target cells.

### Ex vivo NK cell activity assays

For degranulation assays using mouse NK cells, total intrahepatic leukocytes or splenocytes were cultured for 4h at 10^7^ cells/mL in RPMI 1640 medium supplemented with 10% FCS, 25 mM HEPES, 1 mM sodium pyruvate, 50 μM 2-ME, MEM nonessential amino acids, penicillin, and streptomycin (all Life Technologies brand; Thermo Fisher Scientific, Waltham, MA, USA). Brefeldin A (1 μg/mL; Sigma-Aldrich) and anti-CD107a (1:100, eBioscience, San Diego, CA) were added to all conditions. For cytotoxicity assays, 15,000 sorted NK cells were cultured with 75,000 YAC-1 target cells in 100 μL RPMI supplemented as before for 24h. Cell death was measured using an LDH cytotoxicity assay (Abcam, Cambridge, U.K.), according to the manufacturer’s instructions. Readings were normalized between the medium only (0% cell death) and lysis buffer (100% cell death) controls.

Degranulation and cytotoxicity assays with human NK cells were carried out as previously described (Cuff et al, 2016). Briefly, freshly isolated intrahepatic lymphocytes were cultured for 4h with Brefeldin A (10 μg/ml; Sigma-Aldrich), Monensin (2 μM; Sigma-Aldrich), and 5 ng/ml PerCP–eFluor 710-conjugated anti-human CD107a (clone eBioH4A3; eBioscience). Cell surface staining was performed at the end of the assay. For cytotoxicity assays, sorted cells were cultured with K562 for 4h at a 1:1 ratio in 50 μl of RPMI 1640 medium supplemented as above. At the end of the assay, target cell death was assessed using Annexin V-FITC (BD Biosciences) and propidium iodide.

### Cell culture experiments

Liver conditioned medium was produced by culturing 500 mg of finely minced liver tissue in serum-free M199 medium (Gibco brand, Thermo Fisher Scientific) for 48h. The medium was cleared of debris by being passed through a 70 μm strainer and by centrifugation at 500 xg for 5 minutes at 4°C. Splenic NK cells were cultured for 24h in RPMI supplemented as before mixed 3:1 with liver conditioned medium or unconditioned M199 medium. TGFβ1 (10 ng/mL, Pepro-Tech, Rocky Hill, NJ), anti-mouse TGFβ (10 μg/mL, clone 1D11 RD Systems, Minneapolis, MN) or isotype control antibody (10 μg/mL, clone 11711 RD Systems) were added as indicated.

### Flow cytometry

Details of antibodies used in the study are given in Figure 8. The lineage cocktail for mouse cells consisted of CD3, CD8α, CD19 and Gr1. Dead cells were excluded using fixable viability dye eFluor 450 (eBioscience) (4°C, 15 minutes). Surface staining was carried out in PBS supplemented with 1% FCS (4°C, 15 minutes). Intracellular staining was carried out using Human FoxP3 Buffer (BD Biosciences, Oxford, UK), except for pS6 staining, which was carried out using Cytofix/cytoperm (BD Biosciences). Data were acquired on an LSRFortessa II (BD Biosciences) and analyzed using FlowJo v.X.0.7 (Tree Star, Ashland, OR, USA). Sorting was carried out on an Aria (BD Biosciences).

**Fig. 8.**
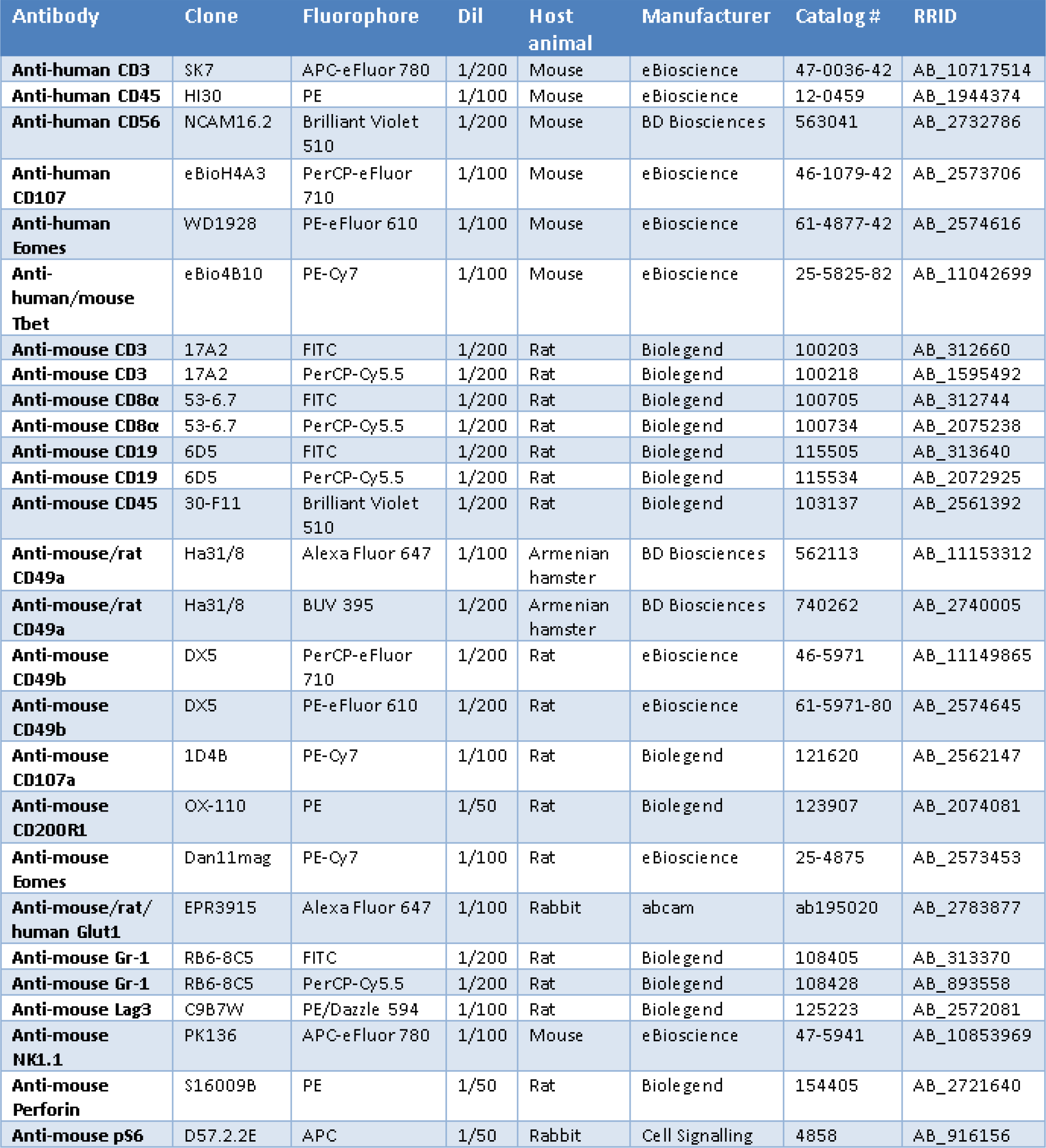
Antibodies used in the study.

### Real time PCR

Liver sections were dissected directly into RNAlater and RNA was extracted using an RNeasy Lipid Tissue Mini kit (both from Qiagen, Manchester, U.K.). cDNA was made using a Transcriptor First Strand cDNA Synthesis kit (Roche, Welwyn Garden City, U.K.) and realtime PCR was performed using TaqMan (Applied Biosystems, Warrington, U.K.) primer/probe sets recognizing Hprt1 (Mm00446968_m1), Acta2 (Mm01546133_m1), Col1a1 (Mm00801666_g1) and Tgfb1 (Mm01178820_m1).

### RNASeq

Total RNA was extracted from sorted cells using an RNeasy Micro kit (Qiagen) and libraries were prepared from 2 ng of total RNA using the NEBNext low input kit (New England Biolabs, Hitchen, U.K.). Libraries were assessed for correct size distribution on the Agilent 2200 TapeStation and quantified by Qubit DNA High Sensitivity assay (Thermo Fisher Scientific) before being pooled at an equimolar concentration. Samples were sequenced on a NextSeq 500 (Illumina, Essex, U.K). Differential expression analysis was carried out using SARTools (Varet et al, 2016), filtering at p_adj_ < 0.05 and FC > 2.

### TGFβ1 ELISA

Total TGFβ1 protein concentrations in plasma and liver conditioned medium were determined using a mouse TGF-beta1 DuoSet ELISA (RD Systems), according to the manufacturer’s instructions.

### Statistics

The data was analyzed and found not to be normally distributed. Significance was therefore determined using non-parametric tests: Mann Whitney U Tests (for unpaired data), Wilcoxon signed rank tests (for paired data) or Spearman’s tests (for correlations) as indicated in the figure legends. p values < 0.05 are reported. Bars represent medians and interquartile ranges; individual data points are also plotted.

## Author Contributions

VM designed the study and secured funding; AOC, FS, SD, AH, TL and VM carried out experiments; AOC and VM acquired data; RC and VM analyzed data; AOC and VM wrote the manuscript; all authors read and approved the manuscript.

## References

Anderson A.C., Joller N., Kuchroo V.K. Lag-3, Tim-3, and TIGIT: Co-inhibitory Receptors with Specialized Functions in Immune Regulation. Immunity. 44:989–1004 10.1016/j.immuni.2016.05.001

Beraza N., Malato Y., Sander L.E., Al-Masaoudi M., Freimuth J., Riethmacher D., et al. 2009. Hepatocyte-specific NEMO deletion promotes NK/NKT cell-and TRAIL-dependent liver damage. J Exp Med. 206:1727–1737 10.1084/jem.20082152

Brunt E.M., Kleiner D.E., Wilson L.A., Belt P., Neuschwander-Tetri B.A.; NASH Clinical Research Network (CRN). 2011. Nonalcoholic fatty liver disease (NAFLD) activity score and the histopathologic diagnosis in NAFLD: distinct clinicopathologic meanings. Hepatology. 53:810–820 10.1002/hep.24127

Castro W., Chelbi S.T., Niogret C., Ramon-Barros C., Welten S.P.M., Osterheld K., et al. 2018. The transcription factor Rfx7 limits metabolism of NK cells and promotes their maintenance and immunity. Nat Immunol. 19:809–820 10.1038/s41590-018-0144-9

Cortez V.S., Ulland T.K., Cervantes-Barragan L., Bando J.K., Robinette M.L., Wang Q., et al. SMAD4 impedes the conversion of NK cells into ILC1-like cells by curtailing non-canonical TGF-β signaling. Nat Immunol. 18:995–1003 10.1038/ni.3809

Cuff A.O., Robertson F.P., Stegmann K.A., Pallett L.J., Maini M.K., Davidson B.R., Male V. 2016. Eomeshi NK Cells in Human Liver Are Long-Lived and Do Not Recirculate but Can Be Replenished from the Circulation. J Immunol. 197:4283–4291 10.4049/jimmunol.1601424

Cuff A.O., Male V. 2017. Conventional NK cells and ILC1 are partially ablated in the livers of Ncr1 iCreTbx21 fl/fl mice. Wellcome Open Res. 2:39 10.12688/wellcomeopen-res.11741.2

Daussy C., Faure F., Mayol K., Viel S., Gasteiger G., Charrier E., et al. 2014. T-bet and Eomes instruct the development of two distinct natural killer cell lineages in the liver and in the bone marrow. J Exp Med. 211:563–577 10.1084/jem.20131560

Dunn C., Brunetto M., Reynolds G., Christophides T., Kennedy P.T., Lampertico P., et al. 2007. Cytokines induced during chronic hepatitis B virus infection promote a pathway for NK cell-mediated liver damage. J Exp Med. 204:667–680 10.1084/jem.20061287

Fehniger T.A., Cai, S.F., Cao X., Bredemeyer A.J., Presti R.M., French A.R., Ley T.J. 2007. Acquisition of murine NK cell cytotoxicity requires the translation of a pre-existing pool of granzyme B and perforin mRNAs. Immunity 26:798–811. 10.1016/j.immuni.2007.04.010

Fernandez I., Zeiser R., Karsunky H., Kambham N., Beilhack A., Soderstrom K., et al. 2007. CD101 surface expression discriminates potency among murine FoxP3+ regulatory T cells. J Immunol. 179:2808–2814 10.4049/jimmunol.179.5.2808

Gao Y., Souza-Fonseca-Guimaraes F., Bald T., Ng S.S., Young A., Ngiow S.F., et al. 2017. Tumor immunoevasion by the conversion of effector NK cells into type 1 innate lymphoid cells. Nat Immunol. 18:1004–1015 10.1038/ni.3800

Hart K.M., Fabre T., Sciurba J.C., Gieseck R.L. 3rd, Borthwick L.A., Vannella K.M., et al. 2017. Type 2 immunity is protective in metabolic disease but exacerbates NAFLD collaboratively with TGF-β. Sci Transl Med. 9:pii: eaal3694 10.1126/scitranslmed.aal3694

Hirsova P., Weng P., Salim W., Bronk S.F., Griffith T.S., Ibrahim S.H., Gores G.J. 2017. TRAIL Deletion Prevents Liver, but Not Adipose Tissue, Inflammation during Murine Diet-Induced Obesity. Hepatol Commun. 1:648–662 10.1002/hep4.1069

Hoek R.M., Ruuls S.R., Murphy C.A., Wright G.J., Goddard R., Zurawski S.M., et al. Down-regulation of the macrophage lineage through interaction with OX2 (CD200). Science. 290:1768–1771 10.1126/science.290.5497.1768

Idrissova L., Malhi H., Werneburg N.W., LeBrasseur N.K., Bronk S.F., Fingas C., et al. TRAIL receptor deletion in mice suppresses the inflammation of nutrient excess. J Hepatol. 62:1156–1163 10.1016/j.jhep.2014.11.033

Kahraman A., Schlattjan M., Kocabayoglu P., Yildiz-Meziletoglu S., Schlensak M., Fingas C.D., et al. 2010. Major histocompatibility complex class I-related chains A and B (MIC A/B): a novel role in nonalcoholic steatohepatitis. Hepatology 51:92–102 10.1002/hep.23253.

Kärre K., Ljunggren H.G., Piontek G., Kiessling R. 1986. Selective rejection of H-2-deficient lymphoma variants suggests alternative immune defence strategy. Nature. 319:675–678 10.1038/319675a0

Lee B.C., Kim M.S., Pae M., Yamamoto Y., Eberlé D., Shimada T., et al. 2016. Adipose Natural Killer Cells Regulate Adipose Tissue Macrophages to Promote Insulin Resistance in Obesity. Cell Metab. 23:685–98. 10.1016/j.cmet.2016.03.002

Luci C., Vieira E., Perchet T., Gual P., Golub R. 2019. Natural Killer Cells and Type 1 Innate Lymphoid Cells Are New Actors in Non-alcoholic Fatty Liver Disease. Front Immunol. 2019 May 28;10:1192. 10.3389/fimmu.2019.01192

Male. V. 2017. Liver-Resident NK Cells: The Human Factor. Trends Immunol. 38:307–309 10.1016/j.it.2017.02.008

Male V., Stegmann K.A., Easom N.J., Maini M.K. 2017. Natural Killer Cells in Liver Disease. Semin Liver Dis. 37:198–209 10.1055/s-0037-1603946

Michelet X., Dyck L., Hogan A., Loftus R.M., Duquette D., Wei K., et al. 2018. Metabolic reprogramming of natural killer cells in obesity limits antitumor responses. Nat Immunol. 19:1330–1340 10.1038/s41590-018-0251-7

Mouralidarane A., Soeda J., Visconti-Pugmire C., Samuelsson A.M., Pombo J., Maragkoudaki X., et al. 2013. Maternal obesity programs offspring nonalcoholic fatty liver disease by innate immune dysfunction in mice. Hepatology. 58:128–138 10.1002/hep.26248

Oben J.A., Mouralidarane A., Samuelsson A.M., Matthews P.J., Morgan M.L., McKee C., et al. 2010. Maternal obesity during pregnancy and lactation programs the development of offspring non-alcoholic fatty liver disease in mice. J Hepatol. 52:913–920 10.1016/j.jhep.2009.12.042

O’Brien K.L., Finlay D.K. 2019. Immunometabolism and natural killer cell responses. Nat Rev Immunol. 19:282–290. 10.1038/s41577-019-0139-2

O’Rourke R.W., Meyer K.A., Neeley C.K., Gaston G.D., Sekhri P., Szumowski M., et al. Systemic NK cell ablation attenuates intra-abdominal adipose tissue macrophage infiltration in murine obesity. Obesity (Silver Spring) 22:2109–14. 10.1002/oby.20823

O’Shea D., Cawood T.J., O’Farrelly C., Lynch L. 2010. Natural killer cells in obesity: impaired function and increased susceptibility to the effects of cigarette smoke. PLoS One. 5:e8660 10.1371/journal.pone.0008660

O’Shea D., Hogan A.E. 2019. Dysregulation of Natural Killer Cells in Obesity. Cancers (Basel). 11:e573. 10.3390/cancers11040573.

Park J., Morley T.S., Kim M., Clegg D.J., Scherer P.E. 2014. Obesity and cancer–mechanisms underlying tumour progression and recurrence. Nat Rev Endocrinol. 10:455–465 10.1038/nrendo.2014.94

Peng, H., Tian, Z. 2015. Re-examining the origin and function of liver-resident NK cells. Trends Immunol. 36:293–299 10.1016/j.it.2015.03.006.

Peng H., Tian Z. 2017. Diversity of tissue-resident NK cells. Semin Immunol. 31:3–10 10.1016/j.smim.2017.07.006

Radaeva S., Sun R., Jaruga B., Nguyen V.T., Tian Z., Gao B. 2006. Natural killer cells ameliorate liver fibrosis by killing activated stellate cells in NKG2D-dependent and tumor necrosis factor-related apoptosis-inducing ligand-dependent manners. Gastroenterology 130:435–452 10.1002/hep.23253.

Snelgrove R.J., Goulding J., Didierlaurent A.M., Lyonga D., Vekaria S., Edwards L., et al. 2008. A critical function for CD200 in lung immune homeostasis and the severity of influenza infection. Nat Immunol. 9:1074–1083 10.1038/ni.1637

Sojka D.K., Plougastel-Douglas B., Yang L., Pak-Wittel M.A., Artyomov M.N., Ivanova Y., et al. 2014. Tissue-resident natural killer (NK) cells are cell lineages distinct from thymic and conventional splenic NK cells. Elife. 3:e01659 10.7554/eLife.01659

Tobin L.M., Mavinkurve M., Carolan E., Kinlen D., O’Brien E.C., Little M.A., et al. 2017. NK cells in childhood obesity are activated, metabolically stressed, and functionally deficient. JCI Insight. 2:pii: 94939 10.1172/jci.insight.94939

Tosello-Trampont A.C., Krueger P., Narayanan S., Landes S.G., Leitinger N., Hahn Y.S. 2016. NKp46(+) natural killer cells attenuate metabolism-induced hepatic fibrosis by regulating macrophage activation in mice. Hepatology 63:799–812 10.1002/hep.28389

Varet H., Brillet-Guéguen L., Coppée J.Y., Dillies M.A. 2016. SARTools: A DESeq2-and EdgeR-Based R Pipeline for Comprehensive Differential Analysis of RNA-Seq Data. PLoS One. 11:e0157022 10.1371/journal.pone.0157022

Viel S., Marçais A., Guimaraes F.S., Loftus R., Rabilloud J., Grau M., et al. 2016. TGF-β inhibits the activation and functions of NK cells by repressing the mTOR pathway. Sci Signal. 9:ra19 10.1126/scisignal.aad1884

Viel S., Besson L., Charrier E., Marçais A., Disse E., Bienvenu J., et al. 2017. Alteration of Natural Killer cell phenotype and function in obese individuals. Clin Immunol. 177:12–17. 10.1016/j.clim.2016.01.007

Weizman O.E., Adams N.M., Schuster I.S., Krishna C., Pritykin Y., Lau C., et al. 2017. ILC1 Confer Early Host Protection at Initial Sites of Viral Infection. Cell. 171:795–808 10.1016/j.cell.2017.09.052

Wensveen F.M., Jelenčić V., Valentić S., Šestan M., Wensveen T.T., Theurich S., et al. 2015. NK cells link obesity-induced adipose stress to inflammation and insulin resistance. Nat Immunol. 16:376–85. doi:10.1038/ni.3120.

Yadav H., Quijano C., Kamaraju A.K., Gavrilova O., Malek R., Chen W., et al. 2011. Protection from obesity and diabetes by blockade of TGF-β/Smad3 signaling. Cell Metab. 14:67–79 10.1016/j.cmet.2011.04.013

Zaiatz-Bittencourt V., Finlay D.K., Gardiner C.M. 2018. Canonical TGF-β Signaling Pathway Represses Human NK Cell Metabolism. J Immunol. 200:3934–3941 10.4049/jimmunol.1701461.

